# SARS-CoV-2 E and 3a proteins are inducers of pannexin currents

**DOI:** 10.1101/2022.10.20.513002

**Authors:** Barbara Oliveira Mendes, Malak Alameh, Béatrice Ollivier, Jérôme Montnach, Nicolas Bidère, Frédérique Souazé, Nicolas Escriou, Flavien Charpentier, Isabelle Baró, Michel De Waard, Gildas Loussouarn

## Abstract

Controversial reports have suggested that SARS-CoV E and 3a proteins may be viroporins that conduct currents through the plasma membrane of the infected cells. If true, these proteins would represent accessible targets for the development of new antiviral drugs by using high-throughput patch-clamp techniques. Here we aimed at better characterizing the cell responses induced by E or 3a protein with a particular focus on the ion conductances measured at the cell surface. First, we show that expression of SARS-CoV-2 E or 3a protein in CHO cells gives rise to cells with newly-acquired round shape, tending to detach from the Petri dish. This suggests that cell death is induced upon expression of E or 3a protein. We confirmed this hypothesis by using flow cytometry, in agreement with earlier reports on other cell types. In adhering cells expressing E or 3a protein, whole-cell currents were in fact not different from the control condition indicating that E and 3a proteins are not plasma membrane viroporins. In contrast, recording currents on detached cells uncovered outwardly-rectifying currents, much larger than those observed in control. The current characteristics are reminiscent of what was previously observed in cells expressing SARS-CoV-1 E or 3a proteins. Herein, we illustrate for the first time that carbenoxolone blocks these outward currents suggesting that they are conducted by pannexin channels, mostly likely activated by cell morphology change and/or cell death. Alongside we also demonstrate that truncation of the C-terminal PDZ binding motifs reduces the proportion of dying cells but does not prevent pannexin currents suggesting distinct pathways for cell death and pannexin currents induced by E and 3a proteins. We conclude that SARS-CoV-2 E and 3a proteins are not acting as viroporins expressed at the plasma membrane.

**Author Summary:** A viroporin (or viral porin) is a class of proteins that is encoded by a virus genome. It is named porin because its biological role is to conduct ions through a pore that it created in a lipid membrane such as the one surrounding a human cell. if such viroporin is present at the external membrane of a human cell infected by a virus, it can be an easy target of an antiviral agent which thus does not have to enter the cell to be active. One example of viroporin is the flu M2 protein that is the target of amantadine, an antiviral agent used against flu. In previous studies, two proteins of SARS-CoV viruses, named E protein and 3a protein, have been suggested to be viroporins at the surface of infected human cells, potentially opening a new research avenue against SARS. Here we demonstrate that both proteins are not viroporins at the external membrane but they rather trigger changes in the cell shape and promote cell death. They only indirectly induce the activity of a porin that is encoded by the cell genome, named pannexin.

## Introduction

SARS-CoV-2 is the third virus of the genus Beta-coronavirus of the Coronaviridae family to be responsible for a Severe Acute Respiratory Syndrome in this century, after SARS-CoV-1 in 2002-2003 (1) and MERS-CoV in 2012 (2). As a result, it is of great importance to best characterize coronaviruses and the associated pathophysiology, with the hope that new treatments will emerge to complement vaccine approaches for people who cannot access the vaccines or are not responsive to them. In addition to paxlovid which is already available but associated with bothersome side-effects (3), many potential anti-COVID-19 treatments are in development but it is too soon to tell how efficient they will be, namely with regard to the continuous emergence of new variants, and if the cost will be reasonable (4).

Viroporins, *i.e.* ion channels encoded by a virus genome, are potential targets by antiviral agents, as demonstrated by the case of amantadine which inhibits the acid-activated M2 channel of Influenza A virus (5). Several studies led to the suggestion that two proteins of SARS-CoV are viroporins. SARS-CoV-2 Envelop (E) protein, is a one-transmembrane-domain membrane protein (75 amino-acids) almost identical to SARS-CoV-1 Envelop protein (95% identity). The SARS-CoV-2 ORF3a (3a) protein is a larger three-transmembrane-domain membrane protein (275 amino-acids) relatively similar to the SARS-CoV-1 3a protein (73% identity).

Regarding the ion channel function of these proteins, there are clearly several contradicting studies: some of them raising intriguing issues and others not confirming these reports. Concerning *in-vitro* membrane incorporation of purified E or 3a protein in lipid bilayers, the presence of ion channel activity was reportedly associated with these viral proteins (6–11). However, a review article soundly outlined the lack of robust data and raised ethical concerns casting doubts on the validity of the scientific messages (12). Concerning viral protein expression in cells, the expression of SARS-CoV-1 E protein also led to conflicting results (13,14). Pervushin et al managed to identify plasma membrane currents generated by heterologous expression of SARS-CoV-1 E protein in HEK-293 cells (13) but not Nieto-Torres et al (14). More recently, expression of wild-type (WT) SARS-CoV-2 E protein did not lead to interpretable ionic currents in HEK-293S or *Xenopus laevis* oocytes (15) despite their high homology among SARS-CoV viruses. In an attempt to favor plasma membrane targeting and reveal a putative current, a C-terminal predicted ER retention signal of SARS-CoV-1 E protein was replaced by a Golgi export signal from Kir2.1 channel. Expression of this chimera could then be associated with the generation of a non-rectifying and cation-selective current. This current was thus quite different from the outwardly rectifying current observed by Pervushin and collaborators (15). Furthermore, another study using a membrane targeting sequence, fused to the N-terminus of the SARS-CoV-1 E protein, provided a non-rectifying current that was 100-fold larger than the one observed in the two previous studies (16). This suggests that such modification of either N- or C-terminus are too drastic to faithfully report the actual activity of the native proteins.

SARS-CoV-1 3a protein was also investigated and expression of the WT protein in HEK-293 cells (17) or *Xenopus laevis* oocytes (18–20) was associated with a poorly selective outwardly rectifying current in both models, resembling to the one observed upon expression of the E protein.

To summarize, there is no unequivocal evidence that SARS-CoV E and 3a proteins are viroporins active at the plasma membrane of the host cell. However, it was recently reported that SARS-CoV-2 E and 3a proteins can promote cell death (21,22), on one hand. On the other hand, apoptosis is associated with the increase of an outwardly-rectifying current conducted by pannexins (23–25). This led us to reinvestigate the actual function(s) of SARS-CoV-2 E and 3a proteins in mammalian cells in the frame of the cell toxicity of these proteins.

In this study, CHO cells expressing either CoV-2 E or 3a proteins tended to develop into a round-shaped form with a tendency to detach from the Petri dish, a process exacerbated compared to control conditions. This cell phenotype is consistent with cell death (26,27) and we confirmed by flow cytometry experiments that expression of E or 3a proteins does indeed promote cell death. Transfected cells, still attached to the Petri dish (adhering cells), had unchanged basal currents indicating that E and 3a proteins are unlikely to act as plasma membrane channels. In contrast, recording whole-cell currents on round-shaped and detached cells, we observed large outwardly-rectifying currents only in E or 3a protein-expressing cells but not in control dying cells. This current is reminiscent of those observed in previous publications using HEK-293 cells and oocytes expressing the SARS-CoV-1 proteins (13,18–20). Application of carbenoxolone, a pannexin channel inhibitor, suggests for the first time that these currents are pannexin-mediated conductances, potentially activated in apoptotic cells. In conclusion, both SARS-CoV-2 E and 3a proteins are most likely triggers of endogenous conductance.

## Results

We first focused on native E and 3a proteins. To maximize the chance of observing E and 3a protein-induced ionic currents, we chose to use pIRES plasmids, in which the protein of interest situated in the first cassette is more expressed than the eGFP reporter in the second cassette, thereby guaranteeing expression of a high level of the protein of interest in fluorescent cells (28,29). For the purpose of this study, we also selected CHO rather than HEK-293 cells because they express minimal endogenous currents (30). We compared whole-cell currents recorded during a ramp protocol, in cells transfected either with a control pIRES2-eGFP plasmid (pIRES), or the same plasmid containing the cDNA of the SARS-CoV-2 E protein (pIRES - E) or 3a protein (pIRES - 3a). Unexpectedly, we did not observe any difference in the currents recorded for the SARS-CoV-2 proteins expressing cells compared to control pIRES condition (Figure 1A). However, many cells transfected with either E- or 3a-encoding plasmids were developing altered morphology, shifting from spindle-like cells to more round cells (Figure 1B), similar to what was previously observed in MDCK cells heterologously expressing SARS-CoV-1 E protein (31). Analysis with Fiji tool confirmed an increase in cell roundness (Figure 1C and suppl. Figure 1). In particular, in patch-clamp experiments, some cells were coming off from the dish bottom by losing adhesion. Cell counting indicated that slightly more cells were losing adhesion when E or 3a protein were expressed (3.4 ± 0.6 % in non-transfected cells, 5.2 ± 1.0% in pIRES condition, 6.6 ± 0.7% in pIRES - E, 6.0 ± 1.2% in pIRES - 3a, n=3-5). As classically performed, currents shown in Figure 1A were recorded from adhering cells while non-adhering cells were disregarded in this initial investigation. Noteworthy, in each condition, both spindle-like and round adhering cells were studied (pIRES: 21 spindle-like and 17 round cells; pIRES E: 9 and 13, pIRES 3a: 18 and 11).

**Figure 1:**
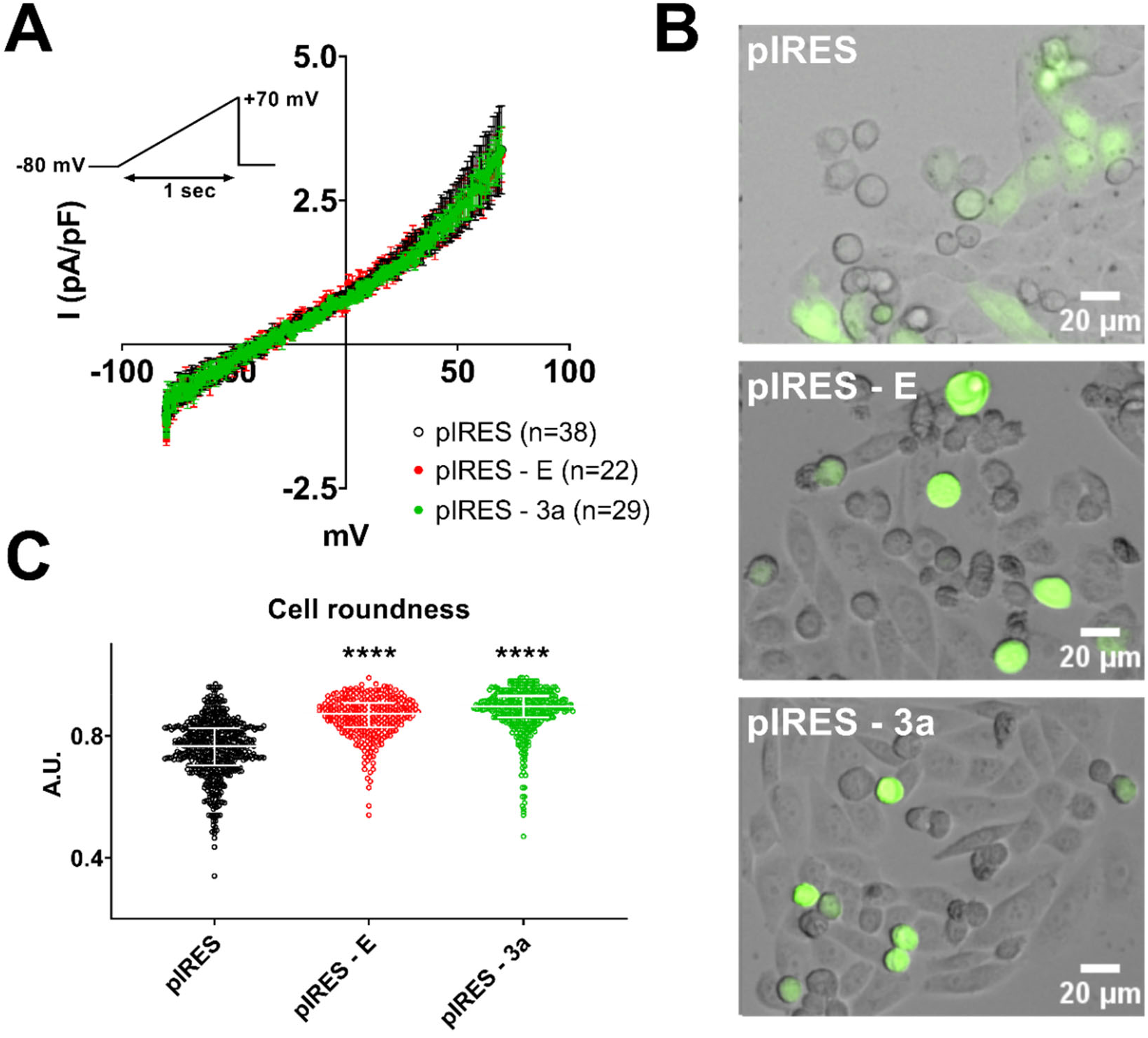
Expression of E and 3a protein is accompanied by altered cellular morphology but no modification in whole-cell currents in adhering cells. **(A)** Average current densities (± sem) recorded during the ramp protocol (inset) in adhering CHO cells expressing either eGFP (pIRES), SARS-CoV-2 E and eGFP proteins (pIRES - E) or SARS-CoV-2 3a and eGFP proteins (pIRES - 3a). **(B)** Superimposed brightfield and eGFP fluorescence images of CHO cells in the same 3 conditions as shown in A. **(C)** Analysis of cell roundness in cell automatically detected by Fiji, cf. suppl. Figure 1. Plot of individual cells (pIRES, n = 1922; pIRES - E, n = 1198; and pIRES - 3a, n = 1283), median ± interquartile range. **** p<0.0001 as compared to pIRES control, Kruskal-Wallis test.

Since both E and 3a proteins are promoting cell death (21,22), we hypothesized that the various cell morphological patterns (spindle-shaped, round-adhering, and round non-adhering) may correspond to the development of cell death, as described earlier in CHO and other cells (26,27). Flow cytometry analysis performed on the eGFP-positive CHO cells (Figure 2) showed that expression of E and 3a proteins increases the percentage of dying cells, more significantly late cell death, revealed by propidium iodide permeability (suppl. Figure 2). The effect of 3a protein was greater than the effect of the E protein. E protein-induced cell death could be reduced by the pan-caspase inhibitor QVD-Oph, while 3a protein-induced cell death was not (suppl. Figures 3 & 4), suggesting that E protein induces apoptosis, while 3a protein activates non-conventional caspase-independent cell death.

**Figure 2:**
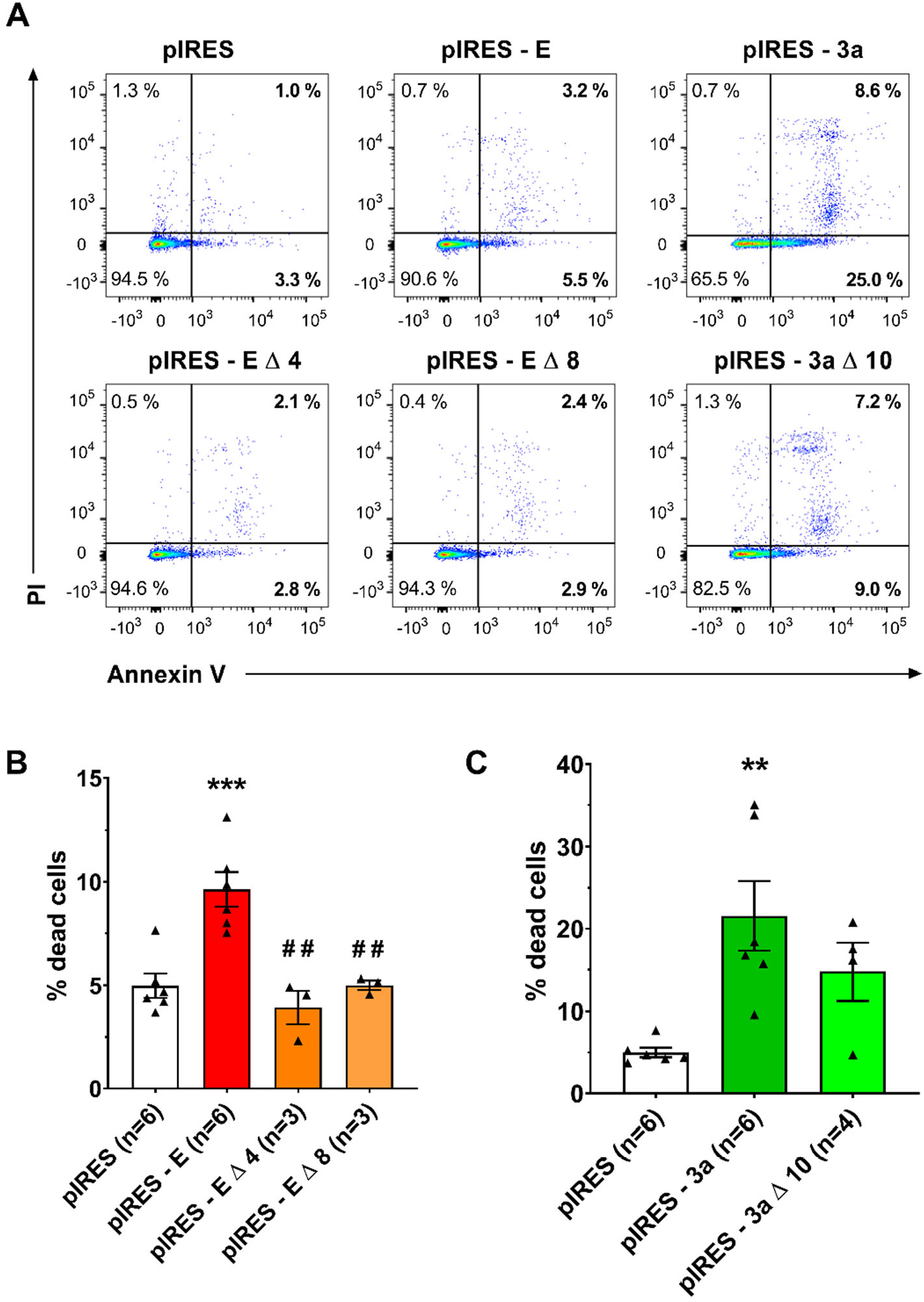
Expression of E and 3a protein induces cell death. **(A)** Flow cytometric analysis of eGFP-positive CHO cells expressing only eGFP (pIRES), or co-expressing eGFP and one of the following SARS-CoV-2 cDNA: the full-length E protein (pIRES - E), C-terminally deleted E protein (pIRES - E Δ 4 or pIRES - E Δ 8), the full length 3a protein (pIRES - 3a) or the C-terminally deleted 3a protein (pIRES - 3a Δ 10). After 48 h of expression, cells were stained with annexin V AlexaFluor 647 (APC)/propidium iodide (PI, Perc-P). **(B)** Mean ± sem of the percentage of Annexin V+ cells among eGFP-positive CHO cells expressing only eGFP (pIRES), or eGFP and full-length or truncated E protein. *** p<0.001 as compared to pIRES control, one-way ANOVA, ## p<0.01 as compared to E protein, t-test. **(C)** Mean ± sem of the percentage of Annexin V+ cells among eGFP-positive CHO cells expressing only eGFP (pIRES), or eGFP and full-length or truncated 3a protein. ** p<0.01 as compared to pIRES control, one-way ANOVA.

Both E and 3a proteins possess a C-terminal PDZ binding motif (PBM). E protein PBM has been suggested to be a virulence factor (11) and binds to host cells PDZ domains, leading to abnormal cellular distribution of the targeted proteins (31). 3a PBM interacts with at least five human PDZ-containing proteins (TJP1, NHERF3 & 4, RGS3, PARD3B), suggesting that it also alters cellular organization (32). We thus evaluated whether deletion of these domains impacts E and 3a proteins propensity to trigger cell death. Two C-terminal deletions used in previous studies to remove E protein PBM, Δ4 for the last 4 amino-acids (31) and Δ8 for the last 8 residues (11) abolished the pro-apoptotic effect of E protein (Figure 2). When looking individually at early and late cell death, we observed that both truncation of E and 3a protein decreased late cell death (suppl. Figure 2).

Since both E and 3a proteins promote cell death, we hypothesized that the cells starting to come off the surface may express currents induced by cell death such as pannexin currents (24,25,33). We thus compared patch-clamp recordings of adhering cells *vs.* non-adhering cells in the 3 conditions: control pIRES, pIRES - E and pIRES - 3a plasmids (Figure 3). In the control pIRES condition, focusing on non-adhering cells in the 35-mm dish and using the ramp protocol, we observed an outwardly rectifying current with a mean current density of 8.5 ± 3.1 pA/pF at +70 mV, slightly higher than in spindle- or round-shaped adhering cells (3.4 ± 0.8 pA/pF). On the other hand, in non-adhering cells expressing either the E or 3a protein, currents were much larger in the E protein condition (I_+70mV_ = 31 ± 9 pA/pF, two-way ANOVA test on the ramp-evoked currents: p<0.0001), but also in the 3a protein condition (I_+70mV_ = 44 ± 13 pA/pF, two-way ANOVA test on the ramp-evoked currents: p<0.0001), as compared to non-adhering cells in the control pIRES condition. Noteworthy, only a fraction of the cells was exhibiting large rectifying currents, as shown in Figure 3: 4 out of 43 in the control pIRES condition, 14 out of 46 in the E protein condition and 16 out of 41 in the 3a protein condition. These experiments suggest that cell death and/or change in morphology induced by expression of E and 3a proteins may lead to an increased membrane permeability by enhancing the expression or activity of an endogenous ion channel.

**Figure 3.**
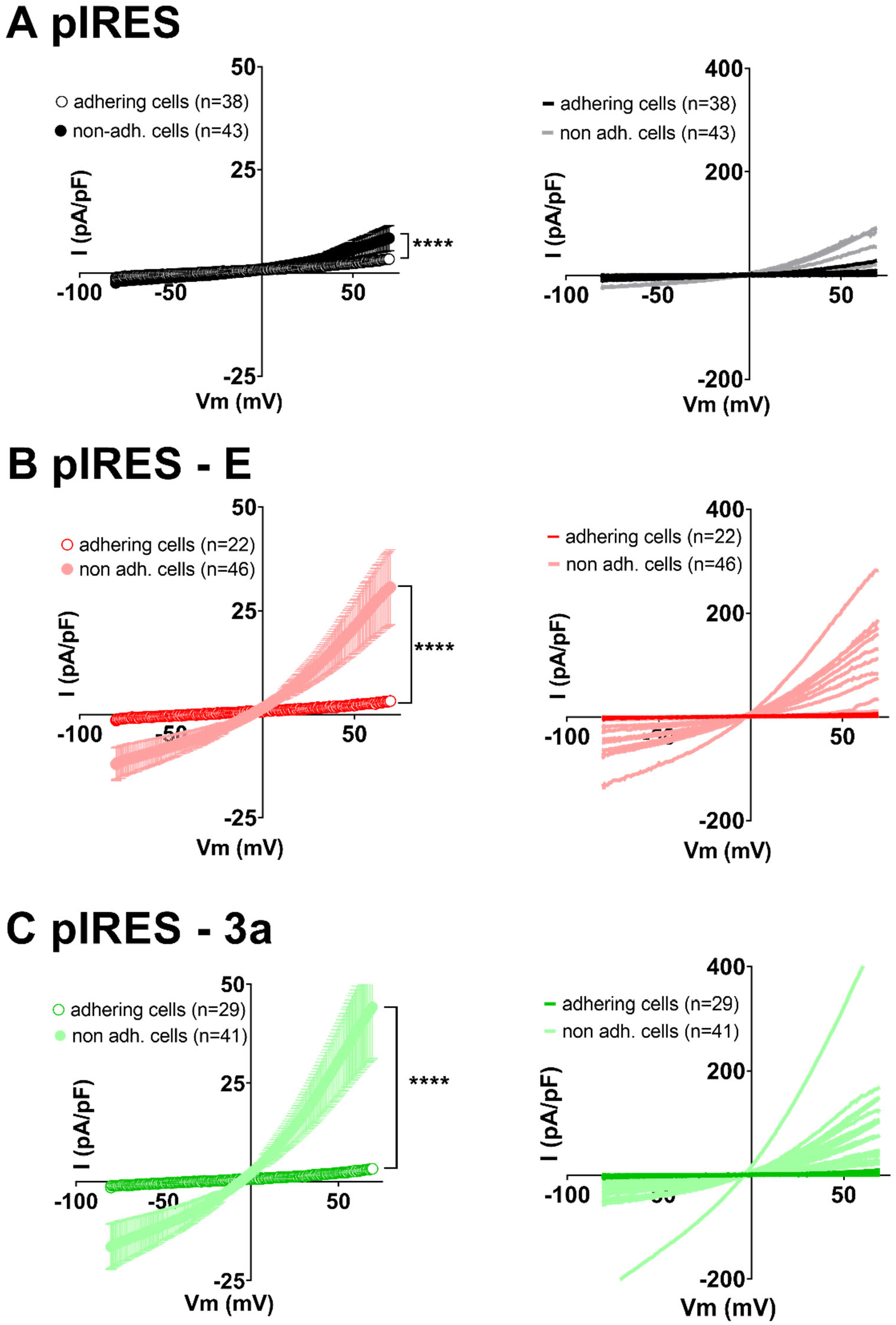
Expression of E and 3a protein is accompanied by outwardly rectifying currents in non-adhering CHO cells only. Left, average current densities (± sem) recorded during the ramp protocol in adhering (empty circles) or non-adhering (filled circles) CHO cells expressing either eGFP (**A**, pIRES), SARS-CoV-2 E and eGFP proteins (**B**, pIRES - E) or SARS-CoV-2 3a and eGFP proteins (**C**, pIRES - 3a). Right, plot of the individual adhering cells (darker color) or non-adhering cells (lighter color). **** p<0.0001, as compared to adhering cells, two-way ANOVA.

The outwardly rectifying currents that we observed resemble apoptosis-induced and stretch-induced pannexin currents (23–25). Thus, we applied the pannexin inhibitor carbenoxolone (CBX) on non-adhering cells that display large outwardly rectifying currents (Figure 4). We observed that CBX, applied at 50 μmol/L, inhibits the observed current, restoring current amplitudes similar to the ones observed in the control cells. Altogether, these observations suggest that the current triggered by the expression of E and 3a protein is a pannexin current.

**Figure 4.**
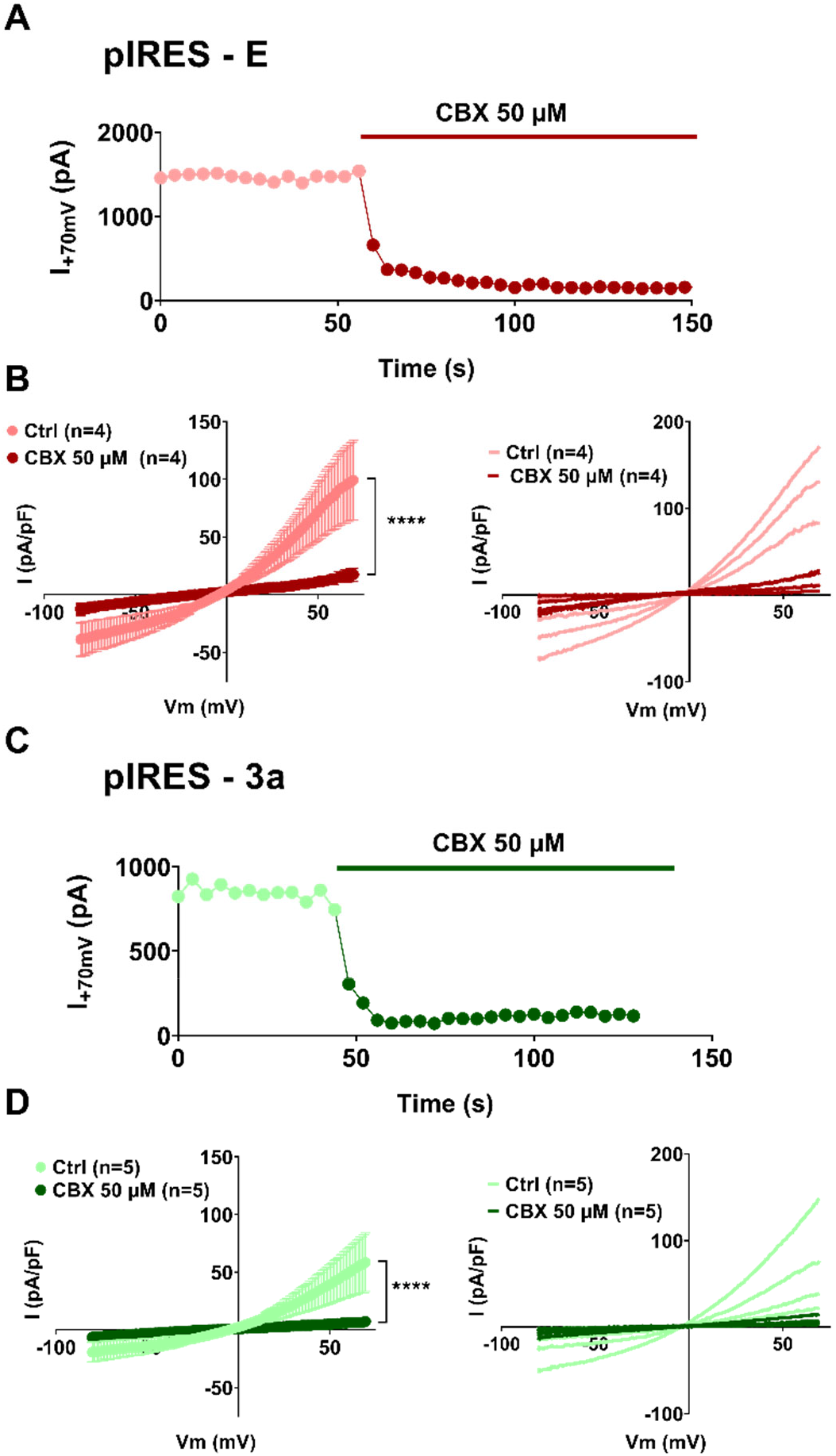
Currents due to E and 3a protein expression in non-adhering CHO cells are suppressed by the pannexin inhibitor carbenoxolone (CBX). **(A)** Recording of the current amplitude in a CHO cell expressing SARS-CoV-2 E and eGFP proteins that displays large outwardly rectifying currents, in absence and presence of CBX. **(B)** Left, average current densities recorded during the ramp protocol in non-adhering CHO cells, expressing SARS-CoV-2 E and eGFP proteins (pIRES - E), in absence (Ctrl, lighter color) and presence of CBX (darker color). Right, plot of the individual non-adhering cells in absence (lighter color) and presence of CBX (darker color). **(C)** Recording of the current amplitude in a CHO cell expressing SARS-CoV-2 3a and eGFP proteins that displays the outwardly rectifying currents, in absence and presence of CBX. **(D)** Left, average current densities recorded during the ramp protocol in non-adhering CHO cells, expressing SARS-CoV-2 3a and eGFP proteins (pIRES - E), in absence (Ctrl, lighter color) and presence of CBX (darker color). Right, plot of the individual non-adhering cells in absence (lighter color) and presence of CBX (darker color). *** p<0.001 or **** p<0.0001, as compared to Ctrl, two-way ANOVA with repeated measures.

We reported above (Figure 2) that deleting the last 4 amino-acids of E protein (Δ4) drastically reduced its pro-apoptotic effect. Cells expressing the Δ4 E protein showed an average roundness similar to cells expressing the WT protein, suggesting that deletion did not prevent its effect on cell morphology (Figure 5A and suppl. Figure 5). Also, when focusing on round and non-adhering Δ4 E protein expressing cells, we could still record large outwardly rectifying currents (5 out of 20 cells), suggesting that C-terminal deletion of E protein does not abolish the induction of pannexin currents, despite prevention of apoptosis probed by flow cytometry experiments (Figure 5B).

**Figure 5.**
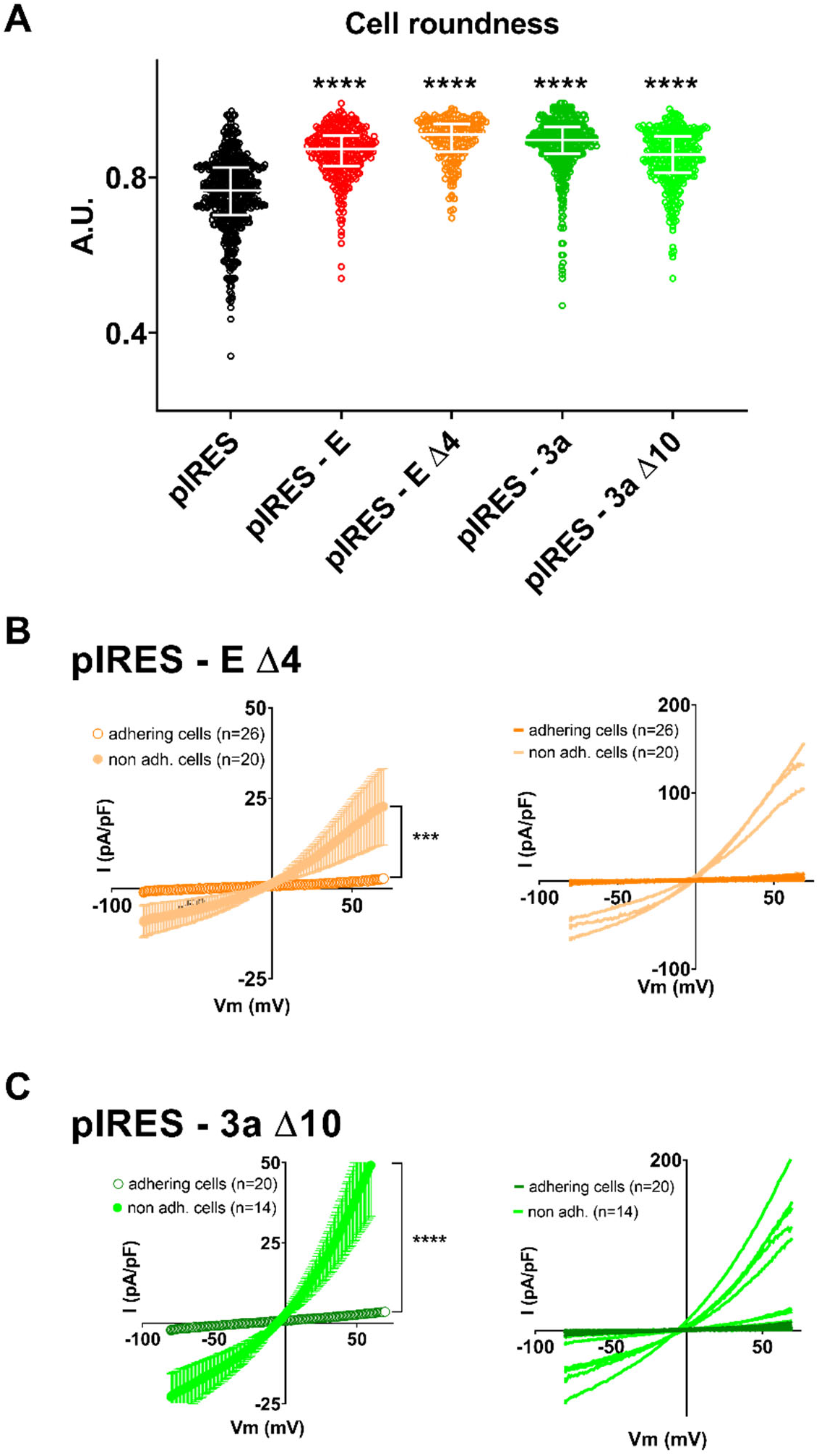
C-terminal deletion of E and 3a protein does not prevent cell characteristic changes. **(A)** Cell roundness measured in CHO cells expressing eGFP (pIRES) alone, or in combination with WT (pIRES - E) or the C-terminally deleted E protein (pIRES - E Δ4) or in combination with WT (pIRES - 3a) or the C-terminally deleted 3a protein (pIRES - 3a Δ10). Plot of individual cells, median ± interquartile range. **** p<0.0001 as compared to pIRES control, Kruskal-Wallis test. **(B)** Left, average current densities (± sem) recorded during the ramp protocol in adhering (empty circles) or non-adhering (filed circles) CHO cells expressing the C-terminally deleted E protein (pIRES - E Δ4). Right, plot of the individual adhering (darker color) or non-adhering cells (lighter color). *** p<0.001, as compared to adhering cells, two-way ANOVA. **(C)** Left, average current densities (± sem) recorded during the ramp protocol in adhering (empty circles) or non-adhering (filled circles) CHO cells expressing the C-terminally deleted 3a protein (pIRES - 3a Δ10). Right, plot of the individual adhering (darker color) or non-adhering cells (lighter color). **** p<0.0001, as compared to adhering cells, two-way ANOVA.

We also reported in Figure 2 that the deletion of the last 10 amino-acids of 3a protein (Δ10) also decreased cell death albeit to a lesser extent. As for E protein deletion, cells expressing the truncated 3a protein showed an average roundness similar to cells expressing the WT protein (Figure 5A and suppl. Figure 5). Focusing on the non-adhering cells, we could still record large outwardly rectifying currents (9 out of 14 cells), suggesting that deletion of the last 10 amino-acids is not sufficient to completely abolish the induction of pannexin-like currents (Figure 5C).

## Discussion

The concept that SARS-CoV E or 3a proteins could be viroporins expressed at the plasma membrane is a seducing one as it could help the identification of new therapeutic drugs against COVID-19 by setting up a screening program on the channel activity. However, this concept was controversial and part of the reasons that explain the controversy about the function of E and 3a proteins is likely linked to the fact that these proteins also trigger morphological alterations and cell death. Cell death is associated to important changes in lipid composition and membrane permeability which suffices to fuel the controversy. One could imagine for instance that cell death would be a way to activate the function of E and 3a proteins at the plasma membrane, but an alternative hypothesis could be simply that cell death triggers an endogenous cell conductance unrelated to the cell function of E and 3a viral proteins. The only way to address these issues was to confirm the cell toxicity, to characterize the impact of the expression of these proteins on cell shape and behavior, to measure plasma membrane conductance, and to characterize them to get an idea of their nature and probe a pharmacological agent that would match the conductance identity. We manage to solve these issues by carefully characterizing the membrane conductances triggered by both E and 3a proteins. The fact that both proteins trigger the same conductance independently of each other was a first indication that they could not be viroporins at the plasma membrane. The second hint was that adhering cells, whether they had a round-shape or not, did not exhibit any outward conductance in spite of E or 3a protein expression. Finally, the sensitivity to carbenoxolone of the outward currents triggered by E or 3a proteins in non-adhering cells was an indication that these viral proteins trigger cellular alterations, such as morphological changes and cell death that are inducers of pannexin-like current. Globally, these observations remain consistent with previous observations that both E and 3a proteins are mainly localized in intracellular compartments in various cell types (14,34–38). Therefore, it is fair to mention that we cannot fully conclude on the viroporin nature of these viral proteins as their localization in subcellular organelles prevents us for clearly testing their intrinsic potential for channel activity.

We showed that expression of either of these two proteins in CHO cells induces an increase in cell death, as quantified by flow cytometry experiments. It is likely, although we did not investigate this point in details, that this cell death accompanies the change in cell morphology and Petri dish detachment. As such, our observation that pannexin-like currents are mainly observed in detached round-shaped cells indicates that major cell morphology changes, up to level of surface detachment, is required for the induction of pannexin-like currents. Whatever the exact mechanism, the upregulation of pannexin channels upon cell death was already observed earlier (39). It is thus not so surprising in fact that other reports faced problems reporting and identifying the conductances triggered by E and 3a viral proteins. The conditions for observing them are indeed quite drastic and require examining cells that are in the combined dying and detachment process, something that is not naturally pursued by researchers, especially if one hopes to detect a viroporin activity. To reconcile our data with earlier publications, we noticed that whole-cell currents observed by others after SARS-CoV-1 E or 3a protein expression in HEK 293 cells (13,17) were also very similar to pannexin currents: outwardly rectifying current, with a reversal potential close to 0 mV at physiological ion concentrations indicating a poor ion selectivity, and amplitude of a few 100 pA.

One possibility is that the pannexin-like currents that we observed are due to the classical caspase-induced cleavage of pannexin (40). Intriguingly, deletion of the C-terminal PBM of E protein abolished its pro-apoptotic effect but cell morphology alteration and pannexin-like currents were still present. Regarding the 3a protein, deletion of its PBM domain only decreased and did not completely abolish the promoted cell death, but again cell morphology alteration and pannexin-like currents were preserved. Altogether, these results suggest that cell morphology modification and pannexin induction may be linked and these processes are not necessarily accompanied by cell death. One has to keep in mind that pannexin currents are activated by many stimuli in addition to cell death (40). In particular, pannexin currents are also stretch-activated and may be enhanced in the detached cells which are undergoing major morphological alterations (23). If pannexins are already activated by stretch, they would not be overactivated by their cleavage by caspase, which would explain the fact that E and 3a proteins truncation do no prevent pannexin current, but only cell death.

In conclusion, SARS-CoV-2 native E and 3a proteins, and probably SARS-CoV-1 ones, do not act as plasma membrane ion channels, but trigger the activity of plasma membrane pannexin channels, most likely through morphological alteration of the cells. However, our study does not rule out potential channel activity in intracellular membranes leading to cell death. Future studies will give more insights on the role of pannexin channels in COVID-19 physiopathology and treatment (41–43).

## Materials and Methods

### Cell culture

The Chinese Hamster Ovary cell line, CHO, was obtained from the American Type Culture Collection (CRL-11965) and cultured in Dulbecco’s modified Eagle’s medium (Gibco 41966-029, USA) supplemented with 10% fetal calf serum (Eurobio, EU), 2 mM L-Glutamine and antibiotics (100 U/mL penicillin and 100 μg/mL streptomycin, Corning, EU) at 5% CO_2_, maintained at 37°C in a humidified incubator. This cell line was confirmed to be mycoplasma-free (MycoAlert, Lonza).

### Drugs

Carbenoxolone disodium salt was purchased from Sigma and 100 mmol/L stock solution was prepared in H_2_O. Drugs used for flow cytometry experiments were QVD-Oph (#OPH001, R&D Systems, 10 mmol/L stock solution in DMSO), S63-845 (Chemieteck, 10 mmol/L stock solution in DMSO), ABT737 (Selleckchem, 10 mmol/L stock solution in DMSO).

### Construction of E and 3a proteins encoding plasmids

SARS-CoV-2 E and 3a nucleotide sequences, containing a Kozak sequence added right before the ATG (RefSeq NC_045512.2) were synthesized by Eurofins (Ebersberg, EU) and subcloned into the pIRES2 vector with eGFP in the second cassette (Takara Bio Europe, EU). Truncated Δ4 and Δ8 E proteins, as well as Δ10 3a protein constructs, lacking the last 12, 24 and 30 last nucleotides, respectively, were also synthesized by Eurofins. Plasmid cDNAs were systematically re-sequenced by Eurofins after each plasmid in-house midiprep (Qiagen, EU).

### E and 3a cDNA transfection

The Fugene 6 transfection reagent (Promega, WI, USA) was used to transfect WT and mutant E and 3a plasmids for patch-clamp, morphology analysis and flow cytometry experiments according to the manufacturer’s protocol. Cells were cultured in 35-mm dishes and transfected, at a 20% confluence for patch clamp experiments and 50% confluence for flow cytometry assay, with a pIRES plasmid (2 μg DNA) with the first cassette empty or containing wild-type or truncated SARS-CoV-2 E or 3a protein sequence. For morphology analysis, cells were cultured in ibidi μ-Slide 8 well dishes and transfected at a 20% confluence with the same plasmids. In pIRES2-eGFP plasmids, the second cassette (eGFP) is less expressed than the first cassette, guaranteeing expression of high level of the protein of interest in fluorescent cells (28,29).

### Electrophysiology

Two days after transfection, CHO cells were mounted on the stage of an inverted microscope and bathed with a Tyrode solution (in mmol/L: NaCl 145, KCl 4, MgCl_2_ 1, CaCl_2_ 1, HEPES 5, glucose 5, pH adjusted to 7.4 with NaOH) maintained at 22.0 ± 2.0°C. Patch pipettes (tip resistance: 2.0 to 2.5 MΩ) were pulled from soda-lime glass capillaries (Kimble-Chase, USA) with a Sutter P-30 puller (USA). A fluorescent cell was selected by epifluorecence. The pipette was filled with intracellular medium containing (in mmol/L): KCl, 100; Kgluconate, 45; MgCl_2_, 1; EGTA, 5; HEPES, 10; pH adjusted to 7.2 with KOH. Stimulation and data recording were performed with pClamp 10, an A/D converter (Digidata 1440A) and a Multiclamp 700B (all Molecular Devices) or a VE-2 patch-clamp amplifier (Alembic Instruments). Currents were acquired in the whole-cell configuration, low-pass filtered at 10 kHz and recorded at a sampling rate of 50 kHz. First, series of twenty 30-ms steps to −80 mV were applied from holding potential (HP) of alternatively −70 mV and −90 mV to subsequently off-line calculate Cm and Rs values from the recorded currents. Currents were then recorded using a 1-s ramp protocol from −80 mV to +70 mV, every 4 s (cf. Figure 1). Regarding non-adhering cells, we considered them as with large current density when the current density measured at +70 mV was superior to mean + 2 x standard deviation of the current density in adhering cells, in the same condition.

### Cell morphology assay

Cell roundness was estimated using the *Analyze Particle* function of the Fiji software, as described in suppl. Figure 1.

### Flow cytometry assay

Two days after transfection, CHO cells were prepared to cell death detection following the user guide (https://assets.thermofisher.com/TFS-Assets/LSG/manuals/mp13199.pdf), to measure annexin V binding and propidium iodide (PI) intake. The cells were washed with cold PBS, trypsinized, collected by centrifugation and gently resuspended in annexin-binding buffer (V13246, Invitrogen, USA) at 1 × 10^6^ cells/mL. To each 300-μL cell suspension were added 0.5 μL of annexin V AlexaFluor 647 (A23204, Invitrogen) and 1 μL of propidium iodide (PI) at 100 μg/mL (P3566, Invitrogen). CHO cells were incubated at room temperature for 15 minutes in the dark, and then maintained on ice until flow cytometry analysis within one hour.

To study the cell death pathways induced by E and 3a protein expression, non-transfected or transfected CHO cells were treated with inhibitors or inducers of apoptosis (inhibitor: 5 μmol/L QVD-OPh incubated for 48 h; activators: 3 μmol/L S63-845 + 8 μmol/L ABT737 incubated for 3 h).

The cytometer BD FACSCanto (BD Biosciences, USA) was used to sample acquisition. CHO cells transfected with an empty plasmid were used to determine the population to be analyzed. Monolabeled cells were used to establish the photomultiplier voltage of each channel (PMT) and to proceed with fluorescence compensation after the acquisitions. In order to detect cell death, only eGFP-positive CHO cells (FITC) were analyzed to Annexin V AlexaFluor 647 (APC) and PI (Perc-P) labeling. Analyses were performed using FlowJo software.

## Acknowledgements

We thank the Agence Nationale de la Recherche for its financial support to the Région Pays de la Loire (ANR FLASH Covid-19 - CoV2-E-TARGET) and the laboratory of excellence ‘Ion Channels, Science and Therapeutics’ (grant No. ANR-11-LABX-0015). We acknowledge the IBISA MicroPICell facility (Biogenouest), member of the national infrastructure France-Bioimaging supported by the French national research agency (ANR-10-INBS-04). We thank the Cytometry Facility Cytocell from Nantes for expert technical assistance. We thank Hugues Abriel and Jean-Sebastien Rougier (Institute of Biochemistry and Molecular Medicine, university of Bern) for fruitful discussions and support and critical reading of the manuscript.

---

**Suppl. Figure 1:**
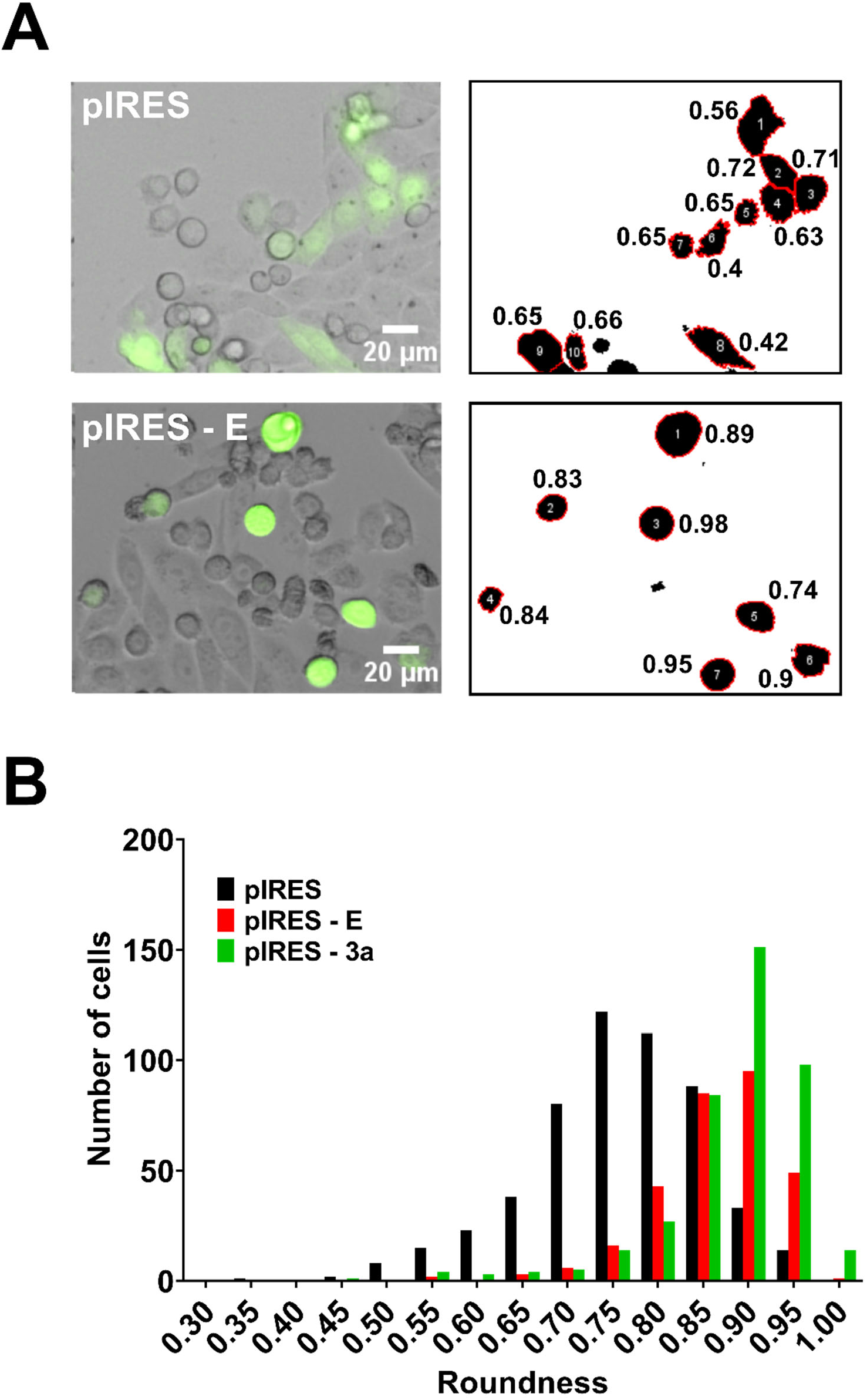
Morphology analysis of cells transfected with control pIRES or pIRES - E plasmids. **(A)** Left: example of superimposed brightfield and eGFP fluorescence images. Right: Automatic particle analysis showing the roundness index **(B)** Distribution of cells roudnesses from the analysis of 5 areas of 1.7 mm x 1.7 mm for cells transfected with control pIRES, pIRES - E or pIRES-3a plasmids.

**Suppl. Figure 2:**
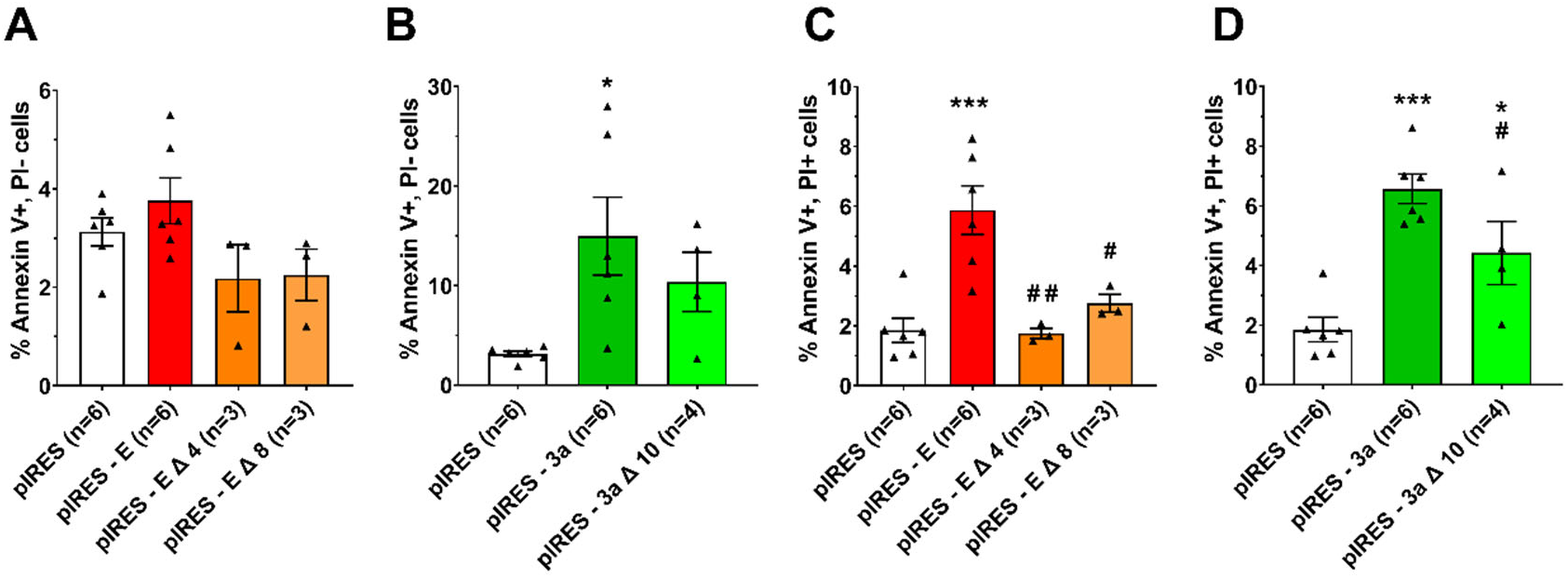
Effects of E and 3a protein expression on early (Annexin V+, PI-in A&B) and late (Annexin V+, PI+ in C&D) cell death. Mean ± sem of the percentage of stained cells among eGFP-positive CHO cells expressing only eGFP (pIRES), or eGFP and full-length or truncated E or 3a protein. * p<0.05, *** p<0.001 as compared to pIRES control, one-way ANOVA, ## p<0.01, # p<0.05 as compared to full length protein, t-test.

**Suppl. Figure 3:**
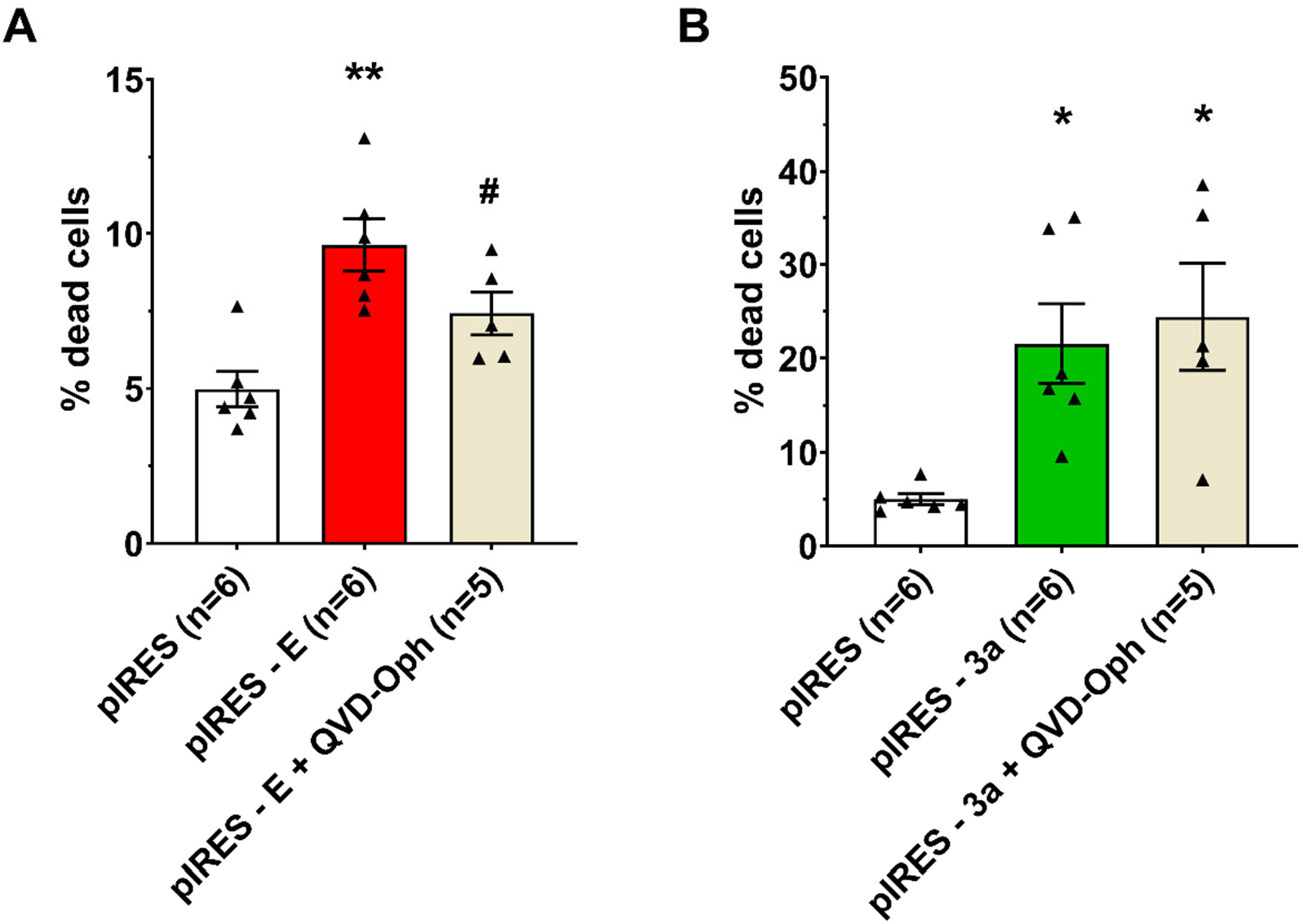
Caspase dependence of E and 3a protein-induced cell death. (**A**) Cell death induced by E protein expression is reduced by the pan-caspase inhibitor QVD-Oph. ** p<0.01, as compared to pIRES control, one-way ANOVA. # p<0.05 as compared to E protein, t-test. (**B**) Cell death induced by 3a protein expression is not reduced by the pan-caspase inhibitor QVD-Oph. * p<0.05, as compared to pIRES control, one-way ANOVA. Concentrations are indicated in the method section.

**Suppl. Figure 4:**
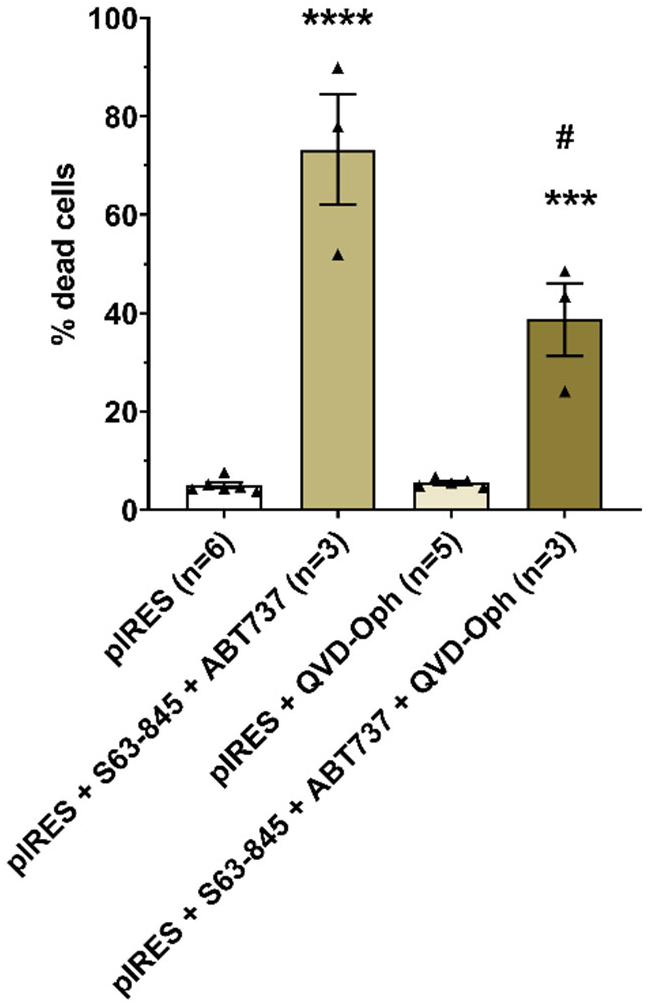
Test of the apoptosis inducers and inhibitor in CHO cells. **** p<0.0001, *** p<0.001, as compared to pIRES control, one-way ANOVA, # p<0.05 as compared to pIRES+S63-845+ABT737, t-test.

**Suppl. Figure 5.**
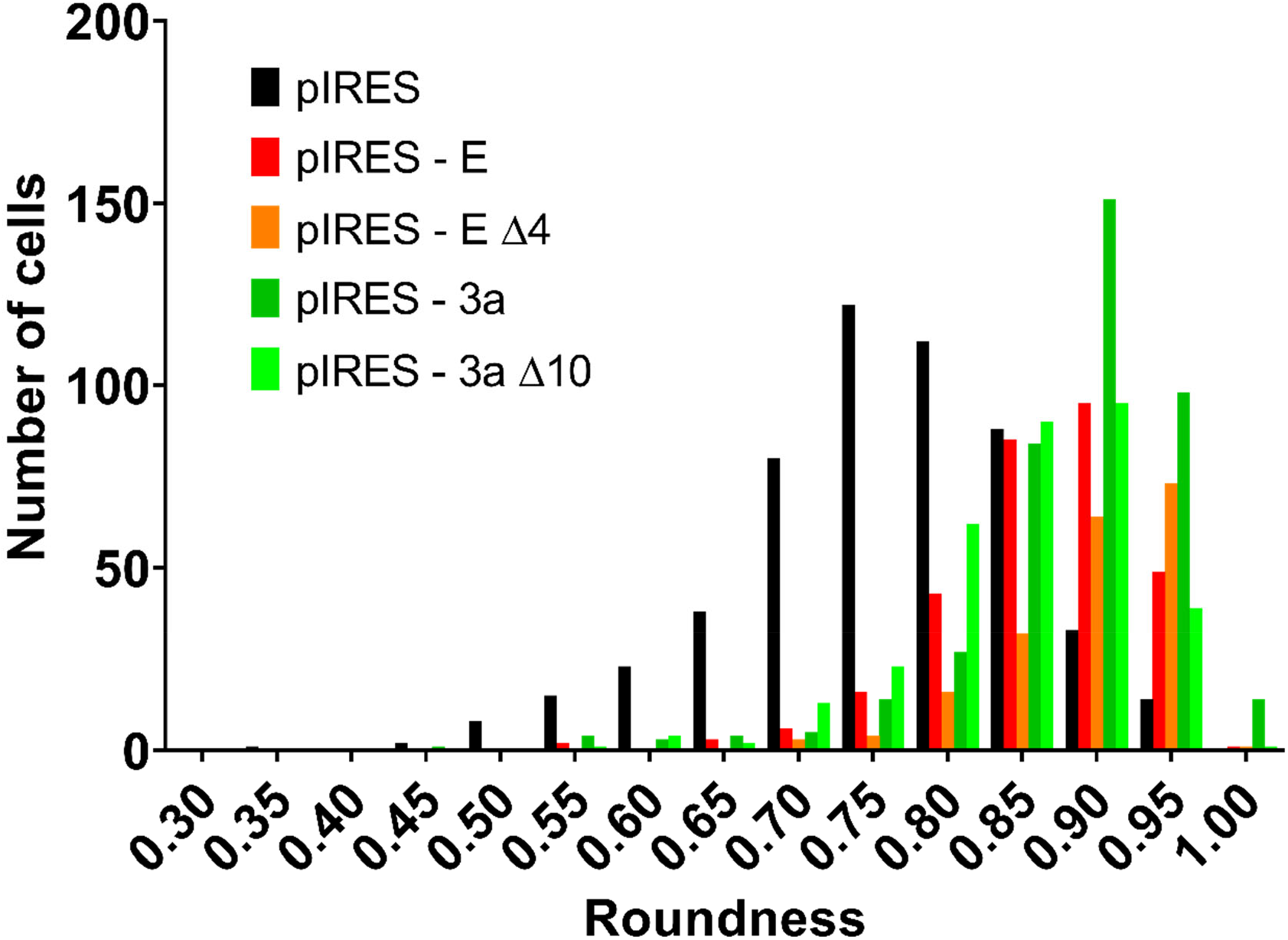
Distribution of cells roundness from the analysis of 5 areas of 1.7 mm x 1.7 mm of transfected CHO cells (Cf. Figure 5A).

